# Uncoupling Drug and Microbiota Contributions to Chemotherapy-Induced Gut Toxicity

**DOI:** 10.64898/2026.06.30.735444

**Authors:** Hadar Bootz-Maoz, Sara Del Mare-Roumani, Gitali Naim, Ofir Azriel, Adva Cohen, Yifat Bennet, Efrat Sharon, Sivan Amidror, Nissan Yissachar

**Affiliations:** The Goodman Faculty of Life Sciences, Bar-Ilan University, Ramat-Gan, 5290002, Israel; Bar-Ilan Institute of Nanotechnology and Advanced Materials, Bar-Ilan University, Ramat-Gan, 5290002, Israel; The Leslie and Susan Gonda Multidisciplinary Brain Research Center, Bar-Ilan University, Ramat Gan 5290002, Israel

## Abstract

Cytotoxic chemotherapy remains a cornerstone of cancer treatment but is frequently limited by gastrointestinal toxicity associated with epithelial barrier disruption. Although chemotherapy profoundly perturbs the gut microbiota, it remains unclear whether these microbial alterations actively contribute to intestinal injury or merely reflect collateral tissue damage. Here, we dissect the respective contributions of direct drug toxicity and chemotherapy-induced dysbiosis to gut pathology using cytarabine (Ara-C), a chemotherapeutic agent associated with severe intestinal complications. We show that both Ara-C and post-Ara-C microbiota independently compromise intestinal barrier integrity. However, these effects arise through distinct host transcriptional programs: Whereas Ara-C directly induces interferon-associated inflammatory responses, post-Ara-C microbiota preferentially activate mucosal barrier defense pathways. Together, these findings identify chemotherapy-induced dysbiosis as an active driver of mucosal injury and provide a mechanistic rationale for microbiome-targeted strategies to mitigate treatment-associated toxicity.

## Introduction

Chemotherapy remains a cornerstone of cancer treatment, but its efficacy is fundamentally limited by off-target toxicities. The gastrointestinal (GI) tract is especially sensitive to antiproliferative drugs due to the high turnover rate of its epithelial lining. Indeed, GI toxicity is among the most frequent and severe adverse effects with nearly all patients receiving intensive chemotherapy developing some degree of mucositis, characterized by mucosal inflammation, ulceration, and loss of barrier functions^1^. Clinically, this manifests as abdominal pain, diarrhea, and malabsorption. Additionally, disruption of the epithelial barrier increases intestinal permeability which can lead to bacterial translocation, bacteremia and sepsis^2,3^. These life-threatening complications often force chemotherapy dose reductions, undermining oncologic outcomes ^4^. Thus, maintaining the integrity of the intestinal epithelial barrier during treatment remains a major unmet challenge in cancer therapy.

Over the past decade, the gut microbiome has emerged as a key modulator of host response to chemotherapy. The dense intestinal microbiota engages in extensive metabolic and immunological cross-talk with the host and can biochemically transform many xenobiotics. Commensal bacteria therefore profoundly influence drug metabolism, therapeutic efficacy, and toxicity^4–6^. Further, differences in microbiome composition have been linked to differential treatment outcomes; for instance, certain gut bacteria (such as *Fusobacterium nucleatum*) can modulate tumor responses and even induce chemoresistance^7,8^. Thus, the gut microbiota functions as a metabolic organ influencing drug bioavailability and toxicity, highlighting the need to account for microbial factors in cancer therapy^5^.

Chemotherapy is not only influenced by the gut microbiome, but also exerts profound effects on microbial composition and function. Cytotoxic agents, often administered in conjunction with broad-spectrum antibiotics in clinical practice, can rapidly deplete microbial diversity and drive the gut ecosystem into a dysbiotic state^9–11^. This disruption is typically marked by the loss of keystone anaerobes and the overgrowth of resilient, and often pathogenic, taxa, creating a permissive niche for microbial invasion^5,8^.

While a healthy microbiota maintains intestinal homeostasis by producing short-chain fatty acids, sustaining the mucus layer, and delivering immunoregulatory signals that preserve epithelial barrier integrity^12,13^, microbial dysbiosis compromises these protective functions. Emerging evidence links chemotherapy-induced microbial shifts with impaired mucosal defense, increased intestinal permeability, and elevated risk of systemic infections^10^. Thus, therapeutic strategies aimed at preserving barrier integrity or modulating the gut microbiota are increasingly recognized as promising approaches to mitigate the collateral damage of cancer treatment^5,14^.

Recent advances have highlighted that chemotherapy-induced dysbiosis is not merely a loss of beneficial taxa, but a functional shift in the microbial metabolome. Depletion of protective metabolites, such as indole-3-propionic acid and specific bile acids, further compromises the epithelial regenerative capacity and the mucus layer’s protective function^11,13,15^. Moreover, in the context of hematologic malignancies, this barrier breakdown occurs concurrently with chemotherapy-induced neutropenia, significantly elevating the risk of life-threatening bloodstream infections derived from the gut microbiota^9,16^.

However, as chemotherapeutic agents simultaneously injure the intestinal epithelium and disrupt gut microbial communities, it remains unclear whether microbiota dysbiosis independently contributes to mucosal pathology, or simply reflects collateral damage from direct drug toxicity. Resolving this question is critical for understanding the mechanisms driving treatment-associated complications and for developing microbiota-targeted interventions to preserve barrier function and enhance treatment tolerance^5,17,18^.

To address this challenge, we focused on cytarabine (Ara-C), a nucleoside analog central to the treatment of acute myeloid leukemia (AML) and other hematologic malignancies ^19^. By impairing DNA synthesis in proliferating cells, Ara-C induces profound myelosuppression, mucosal injury, and intestinal barrier dysfunction, with high-dose Ara-C regimens often leading to mucositis and increased gut permeability.

Leveraging the gut organ culture platform we developed^20,21^, we functionally and transcriptionally dissected early colonic responses to Ara-C and to post-Ara-C dysbiotic microbiota. We show that both Ara-C and its associated microbiota independently impair barrier integrity but elicit distinct transcriptional programs. Our findings establish a non-redundant mechanism of toxicity, in which chemotherapy-induced gut microbiota dysbiosis is an active driver of mucosal pathology, offering a mechanistic rationale for microbiome-based interventions to reduce GI toxicity and enhance treatment tolerance.

## Results

### Ara-C rapidly compromises intestinal epithelial integrity in a dose-dependent manner

To characterize the direct impact of Ara-C on intestinal barrier function, we first performed high-resolution trans-epithelial electrical resistance (TEER) measurements in Caco-2 monolayers. Cells were exposed to increasing concentrations of Ara-C (1, 50, or 100 mM), and TEER values were monitored over 24 hours at 1-minute intervals to assess changes in epithelial barrier integrity^22^. Exposure to 100 mM Ara-C induced a mild but significant reduction in TEER values, indicative of compromised epithelial integrity (**Fig. 1A-B)**. In contrast, 1 mM and 50 mM Ara-C did not affect epithelial barrier resistance, suggesting that a threshold concentration is required to elicit barrier disruption under these *in vitro* conditions.

**Fig. 1.**
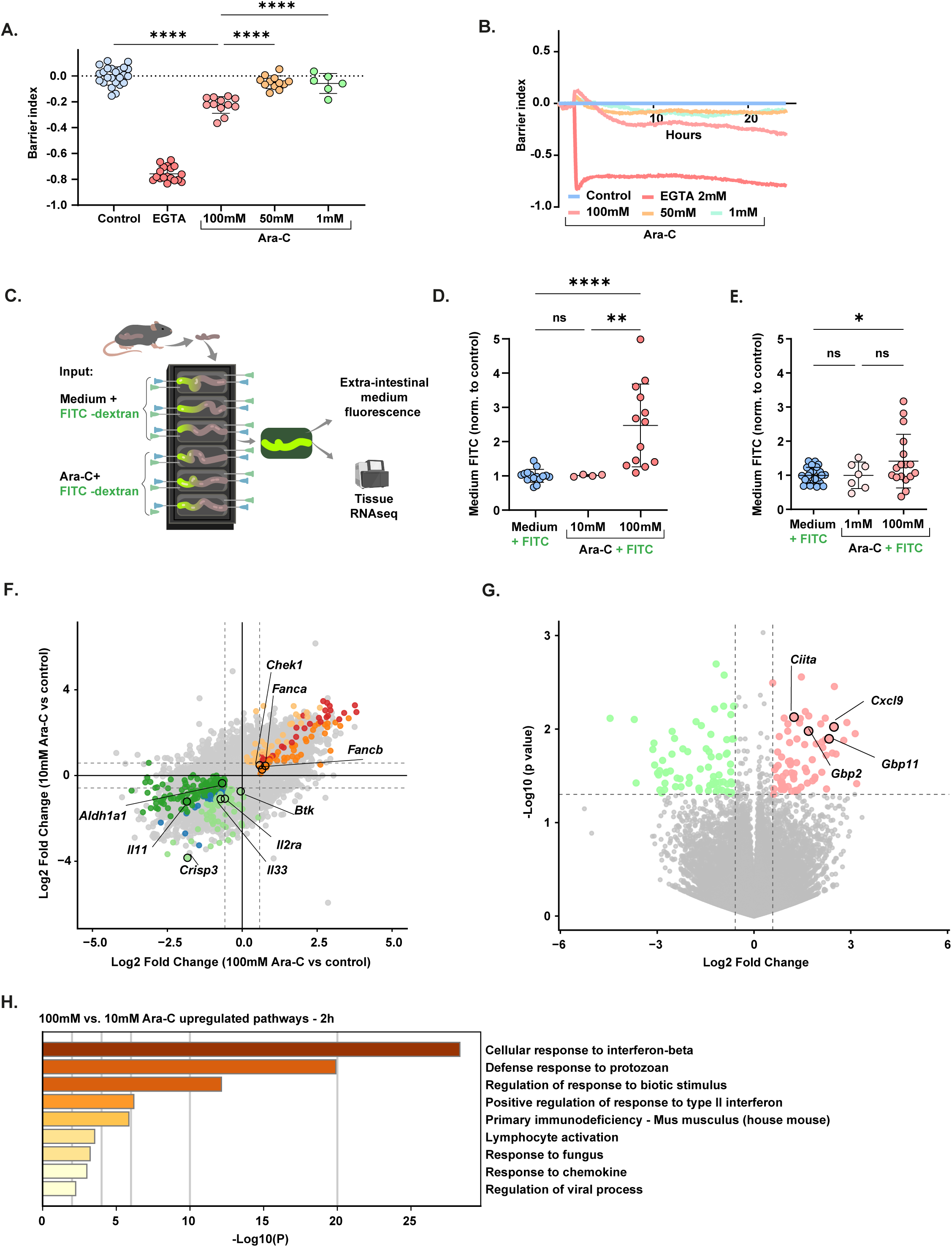
Ara-C directly disrupts epithelial barrier integrity and induces early inflammatory transcriptional responses. **a,b,** Transepithelial electrical resistance (TEER) measurements of differentiated Caco-2 monolayers exposed to Ara-C (1, 50, or 100 mM) or EGTA (2 mM; positive control). **a,** Barrier index at 24 h. **b,** Temporal dynamics of barrier integrity over 24 h, normalized to baseline (t = 0). Data represent pooled measurements from 4 independent experiments (control n = 24, EGTA n = 18, Ara-C 100 mM n = 12, 50 mM n = 12, 1 mM n = 6). **c,** Schematic of the *ex vivo* intestinal permeability assay (X-IPA). Intact mouse colon tissues are mounted in a gut organ culture system and luminally perfused with FITC-dextran (4 kDa) in the presence or absence of Ara-C. Barrier integrity is quantified by fluorescence accumulation in the extraintestinal medium and tissues are collected for downstream transcriptomic analysis. **d,e,** *Ex vivo* permeability of colon tissues following luminal exposure to Ara-C. **d,** FITC-dextran diffusion after 2 h exposure to Ara-C (10 or 100 mM). **e,** FITC-dextran diffusion after 4 h exposure to Ara-C (1 or 100 mM). Fluorescence values are normalized to vehicle-treated controls within each experiment. Each dot represents an individual tissue (2 h: control n = 15, Ara-C 10 mM n = 4, Ara-C 100 mM n = 13; 4 h: control n = 24, Ara-C 1 mM n = 7, Ara-C 100 mM n = 17), pooled from 7 (**d**) or 10 (**e**) independent experiments. **f**, Scatter plot comparing the log₂ fold change (LFC) of gene expression in colon tissues exposed to 100mM Ara-C and 10mM Ara-C treatment, each relative to internal controls (sterile). Each dot represents one gene and is categorized according to its transcriptional response in the two comparisons using *P* ≤ 0.05 and |log₂FC| ≥ 0.58 (abs(FC) >= 1.5). Genes significantly upregulated or downregulated in both conditions are shown in red and blue, respectively. Genes uniquely altered in the 100 mM or 10 mM Ara-C conditions are shown in dark or light orange/green, respectively. Grey dots represent non-significant genes. Dashed lines indicate the applied log2 fold-change thresholds. **g,** Volcano plot showing differential gene expression between 100mM Ara-C and 10mM Ara-C treatment conditions. Significant differences were defined using *P* ≤ 0.05 and |log₂FC| ≥ 0.58. Genes upregulated or downregulated in the 100 mM Ara-C condition are shown in pink and green, respectively. **h,** Metascape pathway enrichment analysis of DEGs in intestinal tissues comparing 100 mM versus 10 mM Ara-C following 2 h stimulation. Significantly upregulated biological pathways are ranked by significance (-log_10_P). **Statistics.** For comparisons across multiple groups, one-way ANOVA with multiple comparisons was used. Data are presented as mean ± s.d. Significance thresholds: \*\*\*\**P* < 0.0001, \*\*\**P* < 0.001, \*\**P* < 0.01, \**P* < 0.05; ns, not significant.

Since Ara-C inhibits DNA synthesis and cell division, we hypothesized that the modest effect observed in slowly dividing Caco-2 monolayers may not fully reflect its barrier-disruptive potential. To further evaluate the direct impact of Ara-C on intestinal permeability, we employed the *ex vivo* intestinal permeability assay (X-IPA) we recently developed ^20^. This system enables quantitative measurement of gut permeability in intact intestinal tissues in response to defined luminal stimuli. Colonic tissues from specific-pathogen-free (SPF) littermate mice were mounted in the X-IPA gut organ culture system and infused with FITC-dextran (4 kDa) in the presence of Ara-C at 1, 10, or 100 mM (**Fig. 1C-E**). Permeability was assessed by measuring fluorescence accumulation in the extraintestinal culture medium at 2 h (**Fig. 1D)** and 4 h (**Fig. 1E**). Exposure to 100 mM Ara-C resulted in a significant increase in permeability at both time-points compared with vehicle-treated sterile controls (**Fig. 1D-E**). Consistent with *the in vitro* TEER measurements (**Fig. 1A-B**), 1 mM and 10 mM Ara-C did not significantly increase permeability at either time point (**Fig. 1D-E**). Together, these results demonstrate that 100mM Ara-C rapidly disrupts epithelial barrier integrity in a concentration-dependent manner both *in vitro* and *ex vivo*.

### Barrier-disruptive Ara-C exposure induces an interferon-associated epithelial injury program

To elucidate the transcriptional programs underlying Ara-C-induced barrier disruption, we performed bulk RNA sequencing on colonic tissues collected 2 h after *ex vivo* exposure to either a barrier-disrupting 100mM Ara-C, a non-disrupting 10mM Ara-C, or untreated control medium (**Fig. 1C**). We have previously shown that this approach captures immediate-early transcriptional responses, providing mechanistic insight into host responses to luminal perturbations^20,21,23–25^.

To visualize the global transcriptional shifts across doses, we compared the log2 fold changes of 100mM and 10mM Ara-C relative to untreated controls (**Fig. 1F, Table S7**). This comparison revealed a broad transcriptional response at 100 mM Ara-C vs. control, with 106 genes significantly upregulated and 201 genes significantly downregulated (abs(fold change) ≥ 1.5, *P*-value ≤ 0.05 (**Table S1**). Among the upregulated genes, we identified a signature of acute genotoxic stress and cell cycle regulation, including *Chek1* (a critical checkpoint kinase activated in response to DNA damage) and components of the Fanconi Anemia DNA repair pathway such as *Fanca* and *Fancb* **(Fig. 1F)**. Additionally, genes involved in mitotic coordination, including *Cenpu* and *Espl1*, were significantly induced, further indicating a robust cellular response to Ara-C-mediated genomic instability (**Fig. S1 B-C**). In parallel, 100mM Ara-C led to the suppression of core epithelial identity markers. This was highlighted by a significant decrease in *Aldh1a1*, a marker of intestinal stemness, signaling hormone *Cck* and *Il11*, which is vital for barrier maintenance and epithelial repair. Consistently, pathway enrichment analysis (Metascape)^26^ revealed enrichment of cell cycle, mitotic progression, and DNA repair pathways among genes upregulated at 100mM Ara-C (**Fig. S1 B**), whereas downregulated genes were associated with epithelial differentiation, and cellular transport (**Fig. S1 C**).

In contrast, comparison of 10mM Ara-C to untreated controls revealed a more restricted transcriptional response with 79 genes significantly upregulated and 91 genes significantly downregulated (abs(fold change) ≥ 1.5, *P*-value ≤ 0.05 (**Fig. S1 D-E Table S2**). As shown in **Fig. 1F**, 10 mM Ara-C selectively suppressed the expression of key immune mediators, including *Btk* (Bruton tyrosine kinase), a central regulator of B-cell function, cytokines and cytokine receptors such as the alarmin *Il33* and *Il2ra*, as well as the pathogen recognition receptor *Clec7a* (Dectin-1) (**Fig. 1F**). Consistently, pathway enrichment analysis indicated that downregulated genes were associated with immune regulatory and stress-response pathways, including modulation of interleukin-4, interleukin-10, and interleukin-17 signaling, regulation of TNF production, response to wounding, and cell-substrate adhesion (**Fig. S1E**). Together, these data indicate that low-dose Ara-C induces a restrained immunomodulatory response, consistent with the established immunosuppressive activity of Ara-C, while lacking the pronounced cell-cycle disruption and tissue injury signatures observed at higher concentrations.

Finally, to define the transcriptional programs associated with the transition from non-disruptive to barrier-disruptive drug exposure, we directly compared tissues treated with 10 mM and 100 mM Ara-C. This analysis identified 74 genes significantly upregulated and 72 genes significantly downregulated in the 100 mM Ara-C condition (abs(fold change) ≥ 1.5, P ≤ 0.05; **Fig. 1G**, **Table S3**). The barrier-disruptive 100 mM Ara-C exposure was characterized by a marked induction of interferon- and immune-associated transcripts, including the interferon-γ-inducible chemokine *Cxcl9*, multiple interferon-stimulated guanylate-binding proteins (*Gbp2, Gbp4, Gbp7*), and regulators of antigen presentation (*Nlrc5* and *Ciita*) (**Fig. 1G**). In agreement, pathway enrichment analysis revealed that the transition to the barrier-disruptive Ara-C dose was dominated by interferon-related pathways, including cellular responses to interferon-β and type II interferon, lymphocyte activation and defense responses to biotic stimuli (**Fig. 1H**). Notably, this interferon-associated transcriptional program was largely absent in the non-disruptive 10 mM Ara-C condition, indicating that robust immune activation emerges specifically at barrier-disruptive Ara-C exposure levels.

Collectively, these data define a direct, dose-dependent trajectory in which 10mM Ara-C induces a limited stress response and immunosuppression, whereas 100mM Ara-C drives epithelial injury and activates an interferon-associated immune program, consistent with (and potentially contributing to) the observed increase in gut permeability and barrier disruption.

### Ara-C increases gut permeability and disrupts microbiome composition in mice

To determine whether Ara-C perturbs intestinal barrier function and host-microbiome interactions *in vivo*, C57BL/6J mice received daily intraperitoneal (i.p.) injections of cytarabine (Ara-C; 3.6 mg per mouse 100mM) or PBS vehicle for four consecutive days (**Fig. 2A**). Ara-C-treated mice exhibited significant weight loss relative to controls (**Fig. 2B-C**), a phenotype consistently observed in both female and male mice (**Fig. S2A-B**). Notably, Ara-C treatment led to increased systemic uptake of orally administered FITC-dextran, indicating compromised epithelial barrier integrity (**Fig. 2D**). In contrast, colon length remained unchanged at this early time point (**Fig. S2C**). To determine whether Ara-C treatment altered gut microbial community composition, we performed paired beta diversity analysis on microbiome samples collected before and after treatment. Bray-Curtis-based principal coordinates analysis (PCoA) revealed a marked treatment-associated shift in microbial composition following Ara-C exposure (**Fig. 2E**).

**Fig. 2.**
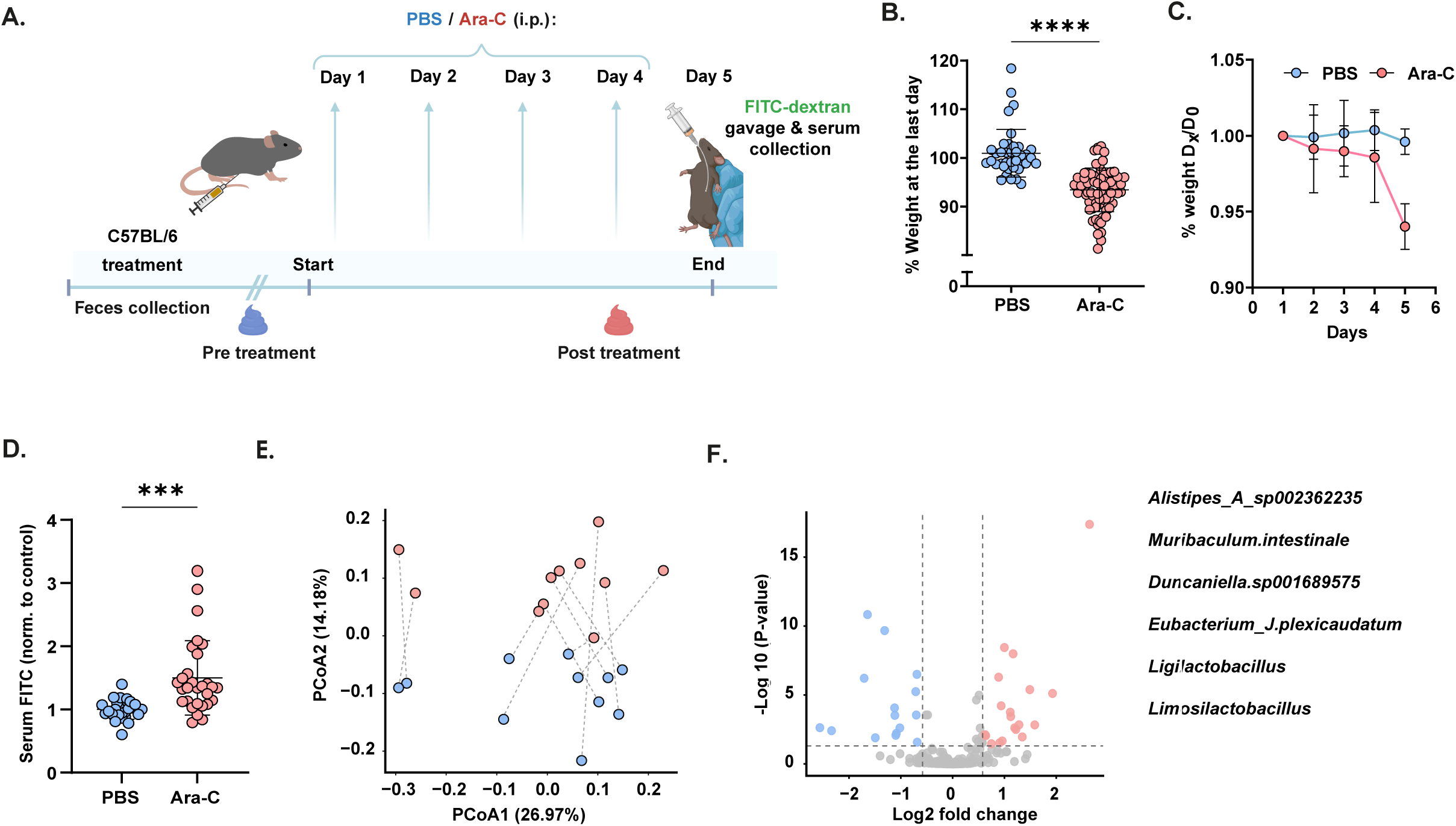
Ara-C treatment induces intestinal barrier dysfunction and microbiota dysbiosis in vivo. **a,** Experimental design of the Ara-C-induced mucositis model. C57BL/6 mice were administered daily i.p. injections of Ara-C or vehicle (PBS) for four consecutive days. Fecal samples were collected longitudinally at baseline (day 0) and throughout treatment. On day 5, following a 12 h fast, mice were orally gavaged with FITC-dextran (4 kDa), and serum fluorescence was quantified 5 h later to assess intestinal permeability. **b,c,** Body weight dynamics during Ara-C treatment. **b,** Final body weight at day 5, expressed as percentage of initial body weight. **c,** Longitudinal changes in body weight over the course of the experiment. Data are pooled from independent experiments (PBS n = 36, Ara-C n = 63; 8 independent experiments). **d,** *In vivo* intestinal permeability assessed by serum FITC-dextran concentration 5 h post-gavage on day 5 (PBS n = 26 blue, Ara-C n = 28 red; pooled from 5 independent experiments). Values are normalized to the mean of the PBS control group. **e,** Bray-Curtis-based Principal Coordinates Analysis (PCoA) of paired microbiome samples collected before and after Ara-C treatment. Each point represents one sample, colored by time point (pre-Ara-C, blue; post-Ara-C, red). Dashed lines connect paired samples from the same mouse to illustrate within-subject temporal microbiome trajectories. For visualization clarity, a representative subset of paired samples is displayed, whereas statistical analysis was performed on the full paired cohort (n=62). (Paired PERMANOVA, P = 0.001). **f,** Volcano plot showing differential abundance of bacterial species between baseline and post-Ara-C microbiota samples. Each dot represents one species plotted by its log₂ fold change versus −log₁₀(*P*-value). Significant differences were defined using *P* ≤ 0.05 and |log₂ fold change| ≥ 0.58. Species enriched or depleted following treatment are shown in red and blue, respectively. **Statistics.** Statistical significance between two groups was assessed using two-tailed unpaired Student’s t-tests (b-d). Data are presented as mean ± s.e.m. Significance thresholds: \*\*\*\**P* < 0.0001, \*\*\**P* < 0.001, \*\**P* < 0.01, \**P* < 0.05; ns, not significant.

Comparative taxonomic profiling between baseline and day 4 identified 34 significantly differentially abundant taxa in the species-level analysis (**Table S4**). Consistent with a dysbiotic shift, Ara-C treatment led to a significant enrichment of several taxa, most notably *Alistipes spp*., (**Fig. 2F)** which have been associated with intestinal inflammation and epithelial barrier dysfunction in both human and murine studies, including in leukemia patients undergoing intensive chemotherapy^27,28^. Conversely, Ara-C treatment resulted in the depletion of multiple taxa essential for gut homeostasis, including members of the *Muribaculaceae* family (such as *Muribaculum intestinale* and *Duncaniella spp.*) and *Eubacterium spp*., as well as several *Lactobacillus*-lineage species (including *Ligilactobacillus* and *Limosilactobacillus*) (**Fig. 2F)**. *Muribaculaceae* taxa are major fermenters of complex polysaccharides that support the production of short-chain fatty acids (SCFAs), which serve as a primary energy source for colonocytes and play a central role in maintaining epithelial barrier integrity and promoting immune tolerance^13,29^. *Eubacterium* spp. include key butyrate-producing bacteria that fuel epithelial metabolism and reinforce tight junction integrity. *Lactobacillus*-lineage commensals further contribute to barrier maintenance by enhancing epithelial junctional complexes, secreting antimicrobial bacteriocins, and suppressing pro-inflammatory signaling pathways. Together, these changes indicate that short-term Ara-C treatment depletes functionally critical barrier-supportive taxa, providing a potential mechanistic link between chemotherapy-induced dysbiosis and impaired epithelial integrity (**Fig. 2F, Table S4**).

### Post-chemotherapy microbiota compromises barrier integrity and triggers pro-inflammatory transcriptional responses

To determine whether chemotherapy-induced dysbiosis contributes to barrier dysfunction, we performed fecal microbiota transplantation (FMT) experiments in which littermate mice were gavaged with fecal microbiota suspensions collected from the same donor mice before or after Ara-C treatment, together with FITC-dextran, thereby enabling paired comparison of pre- and post-treatment microbial communities (**Fig. 3A**). Recipients of post-Ara-C microbiota exhibited a modest but significant increase in serum FITC-dextran levels, indicating elevated intestinal permeability compared with mice receiving pre-treatment microbiota (**Fig. 3B**).

**Fig. 3.**
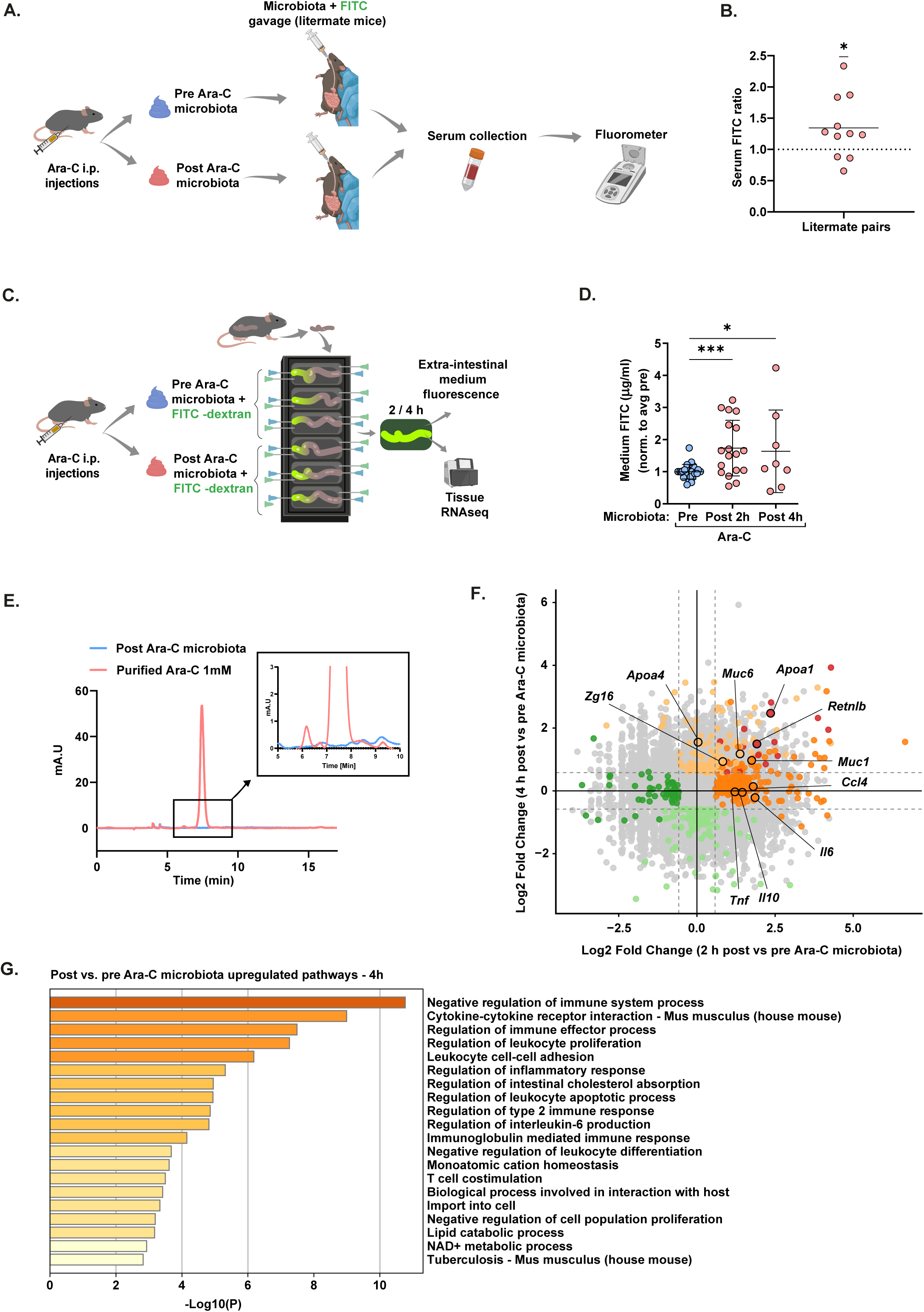
Chemotherapy-altered microbiota is sufficient to induce gut barrier dysfunction *ex vivo* and *in vivo*. **a,** Experimental design for fecal microbiota transplantation (FMT). Littermate C57BL/6 mice were orally gavaged with fecal microbiota collected from donor mice before (pre-Ara-C) or after (post-Ara-C) chemotherapy treatment, as described in Fig. 2a. **b,** *In vivo* intestinal permeability in recipient mice following FMT. Barrier function was assessed by serum FITC-dextran following oral gavage. Each data point represents the ratio of serum FITC levels between littermate pairs receiving post-Ara-C versus pre-Ara-C microbiota from the same donor. Significance was assessed by one-sample t-test (P = 0.0441, n = 22). **c,** Experimental scheme for *ex vivo* intestinal permeability assessment using fecal microbiota. Intact mouse colon tissues were mounted in the gut organ culture system and luminally perfused with medium containing FITC-dextran (4 kDa) together with fecal microbiota derived from pre- or post-Ara-C-treated mice. Extraintestinal fluorescence was quantified at 2 h or 4 h, and tissues were collected for bulk RNA sequencing. **d,** *Ex vivo* intestinal permeability of colon tissues exposed to fecal microbiota. FITC-dextran diffusion into the extraintestinal medium was measured following exposure to microbiota collected before or after Ara-C treatment. Values are normalized to the mean of the pre-Ara-C microbiota condition within each experiment. Each dot represents an individual tissue (pre-Ara-C n = 23, post-Ara-C 2 h n = 13, post-Ara-C 4 h n = 10). **e,** High-performance liquid chromatography (HPLC) analysis of Ara-C. Chromatogram of purified Ara-C (1 mM, red) compared with fecal extracts from Ara-C-treated mice (blue). **f,** Scatter plot comparing the log₂ fold change (LFC) of gene expression in colon tissues 2 h and 4 h following exposure to Ara-C-associated microbiota, each relative to pre-exposure samples. Each dot represents one gene and is categorized according to its transcriptional response in the two comparisons using P ≤ 0.05 and |log₂FC| ≥ 0.58 (abs(FC) >= 1.5). Genes significantly upregulated or downregulated in both conditions are shown in red and blue, respectively. Genes uniquely altered in the 2 h comparison are shown in dark orange/dark green, whereas genes uniquely altered in the 4 h comparison are shown in light orange/light green (upregulated/downregulated, respectively). Selected genes of interest are outlined and labeled. **g,** Metascape enrichment analysis of DEGs in intestinal organ cultures, infused with post-versus pre-Ara-C microbiota following a 4 h stimulation period. Significantly upregulated biological pathways are ranked by significance (-log_10_P). **Statistics.** Statistical analyses were performed using one-way ANOVA with multiple comparisons or one-sample t-test, as indicated in the respective panels. Data are presented as mean ± s.d. Significance thresholds: \*\*\**P* < 0.0001, \*\*\**P* < 0.001, \*\**P* < 0.01, \**P* < 0.05; ns, not significant.

To further assess this effect under controlled conditions, we employed the *ex vivo* intestinal permeability assay (X-IPA ^20^). Intact colon tissues were mounted in the X-IPA organ culture platform and infused with pre- or post-Ara-C microbiota from the same donor, together with FITC-dextran (4 kDa) (**Fig. 3C**). Consistent with the *in vivo* findings, FITC accumulation in the extraintestinal culture medium was significantly increased at both 2 h and 4 h in tissues exposed to post-Ara-C microbiota, indicating compromised epithelial barrier integrity (**Fig. 3D**).

To exclude the possibility that residual Ara-C in fecal samples accounted for these effects, we quantified drug levels by high-performance liquid chromatography (HPLC). Whereas Ara-C was readily detected in the 1 mM spike-in control, it was completely undetectable in fecal samples from Ara-C-treated mice (**Fig. 3E**). These results indicate that post-chemotherapy dysbiotic microbiota impairs epithelial barrier function independently of residual drug exposure.

We next examined the transcriptional consequences of microbiota-driven barrier disruption. RNA sequencing of colonic tissues exposed to pre- or post-Ara-C microbiota revealed a time-dependent transcriptional response, with 272 differentially expressed genes (DEGs) at 2 h and 234 DEG (at 4 h (abs(fold change) ≥ 1.5, *P*-value ≤ 0.05))**Fig. 3F-G, Fig. S3A-C**; **Table S5, Table S6**). A comparison analysis (log2FC 2 h vs. 4 h) identified a core set of genes showing sustained induction across both time points **(Fig. 3F, Table S8)**.

The early response at 2 h was characterized by the induction of classic pro-inflammatory markers and acute stress signals, including the master inflammatory cytokines *Tnf* and *Il6*, alongside the potent chemokines *Ccl2*, *Ccl3*, and *Ccl4*. Other highly induced factors included *Csf3*, the oxidative stress marker *Nos2* (encodes inducible nitric oxide synthase iNOS), and the anti-inflammatory regulator *Il10*. Pathway enrichment analysis further highlighted pathways related to regulation of cell-cell adhesion and extracellular matrix organization, suggesting the early initiation of epithelial remodeling and barrier destabilization programs following Ara-C exposure **(Fig. S3B, Table S5)**.

By 4 h, the transcriptional profile expanded significantly, reflecting a robust activation of innate immunity and barrier defense mechanisms. Upregulated genes included goblet cell-associated factors such as *Muc6* and *Zg16*, consistent with enhanced mucin production and secretory responses, potentially reflecting an early epithelial protective or compensatory program following Ara-C exposure. The 4 h time point was also characterized by a prominent recruitment signal, evidenced by the upregulation of multiple chemokine receptors (*Ccr2*, *Ccr3*, *Ccr5*, *Ccr7*) and the adhesion molecule *Sell* (L-selectin). Furthermore, we identified induction of primary danger signals and metabolic regulators of immunity, including *Ifnb1*, *Card9*, *Ido1*, and *Tnfsf14*, as well as *Ncf4* (which is associated with reactive oxygen species production in phagotyces). Consistent with activation of mucosal defense and repair programs, we also observed induction of the epithelial-derived factor resistin-like molecule beta (*Retnlb*) together with lipid metabolism-associated genes linked to innate lymphoid cell (ILC) biology, including *Apoa1* and *Apoa4* **(Fig. 3F, Table S6)**. Pathway enrichment analysis of differentially expressed genes (**Fig. 3G, Fig. S3A-C**) supported a temporal transition from acute inflammatory signaling toward broader antimicrobial defense and epithelial reinforcement programs. At 4 h, this response was accompanied by coordinated cytokine and immunoregulatory signaling, alongside marked suppression of neuronal and synaptic pathways, suggesting early disruption of neuroepithelial homeostasis following Ara-C exposure (**Fig. 3G, Fig. S3A**).

Together, these findings establish that dysbiotic microbiota from Ara-C-treated mice serve as an independent and sufficient trigger of gut barrier dysfunction and mucosal immune activation, through mechanisms that are distinct from those induced by direct drug toxicity.

### Direct and microbiota-mediated effects of Ara-C elicit distinct colonic transcriptional responses

To distinguish host responses driven by direct chemotherapy toxicity from those induced by chemotherapy-associated dysbiosis, we compared colonic transcriptomes following *ex vivo* exposure to either Ara-C (2 h) (**Fig. 1F-H**) or microbiota isolated from Ara-C-treated mice (2 h and 4 h post-transfer) (**Fig. 3F-G**). Differential expression analysis revealed that the post-chemotherapy microbiota induced a transcriptional response fundamentally distinct from that triggered by direct Ara-C exposure (**Tables S9-S10**). Notably, transcriptional overlap between the two conditions was negligible, with no shared upregulated genes at 2 h (**Fig. 4A**) and only a single shared gene (*S1pr4*) at 4 h (**Fig. 4B**). Together, these findings demonstrate that chemotherapy-induced dysbiosis constitutes an independent driver of intestinal dysfunction, operating through molecular mechanisms distinct from the direct cytotoxic effects of Ara-C.

**Fig. 4.**
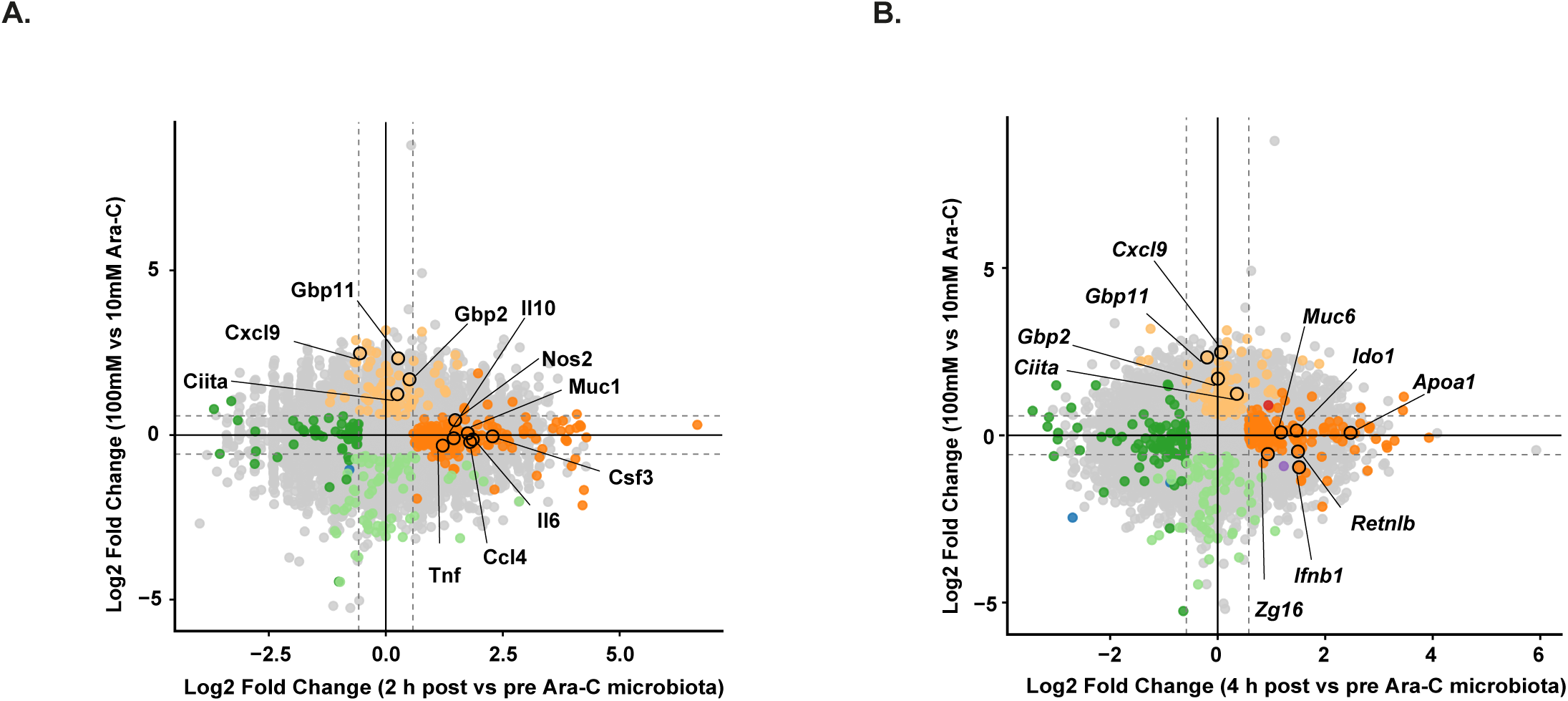
Microbiota-driven barrier dysfunction engages transcriptional programs distinct from direct chemotherapy exposure. **a,b,** Scatter plots comparing log₂ fold changes (LFC) in gene expression between colonic tissues exposed to post-Ara-C versus pre-Ara-C microbiota after **a,** 2 h or **b,** 4 h infusion (x axis), and tissues exposed to 100 mM versus 10 mM Ara-C for 2 h (y axis). Each dot represents one gene and is categorized using P ≤ 0.05 and |log₂FC| ≥ 0.58. Genes significantly upregulated or downregulated in both conditions are shown in red and blue, respectively. Genes uniquely altered in the microbiota comparison are shown in dark orange/dark green, whereas genes uniquely altered in the Ara-C comparison are shown in light orange/light green (upregulated/downregulated, respectively. Selected genes of interest are outlined and labeled.

## Discussion

The intestinal epithelium forms a critical barrier between the host and the external environment and is particularly vulnerable to therapeutic perturbations that compromise barrier integrity. Consequently, chemotherapy-associated gastrointestinal toxicity is generally interpreted as a direct collateral consequence of cytotoxic injury to rapidly proliferating epithelial cells^30^. Our findings indicate that this model is incomplete. We show that cytarabine (Ara-C) not only disrupts intestinal barrier integrity directly, but also generates a dysbiotic gut microbiota that independently impairs barrier function. Importantly, these effects occur through distinct host transcriptional programs and, in the case of the microbiota, in the absence of detectable residual drug. These observations demonstrate that chemotherapy-induced gut toxicity is not solely a host-intrinsic consequence of drug exposure, but also includes a microbiota-mediated component that is mechanistically separable from direct drug injury.

These findings reframe chemotherapy-associated dysbiosis from a secondary consequence of tissue damage to an active driver of pathology. Although cytotoxic therapies are well known to perturb microbial community structure^3,5^, and induce bacterial translocation^31^ it has remained difficult to determine whether these alterations simply mirror epithelial injury or instead contribute causally to treatment-associated complications. By experimentally uncoupling direct drug exposure from microbiota-derived effects using an *ex vivo* gut organ culture platform, we demonstrate that microbiota emerging after chemotherapy is sufficient to impair barrier integrity and initiate host responses distinct from those triggered by Ara-C itself. Thus, part of chemotherapy toxicity appears to be executed by the altered microbial ecosystem that arises following treatment.

These results support a broader framework that can be described as an ecology of toxicity. In this model, an initial insult damages the mucosal barrier and reshapes the microbial ecosystem, generating a dysbiotic community that becomes a secondary effector of pathology. This ecological dimension of toxicity has important therapeutic implications. If part of chemotherapy-associated injury is mediated by dysbiotic microbiota rather than by direct drug action, then this component of toxicity may be therapeutically separable from the anticancer activity of the drug itself. Microbiome-directed interventions including rationally-designed probiotics, synbiotics, dietary modulation, or fecal microbiota transplantation^14,18^ may therefore reduce treatment-limiting toxicity without weakening the intended antitumor mechanism of chemotherapy. In this sense, microbiome modulation should not be viewed solely as supportive care but as a strategy that could widen the therapeutic index of cytotoxic therapy. This perspective is increasingly consistent with emerging oncology literature recognizing host-microbiome interactions as determinants of both treatment efficacy and toxicity ^32^.

More broadly, our findings establish a causality framework for microbiota-driven pathology that may help define clinical contexts in which microbiome-targeted interventions are biologically justified. Such contexts are likely to arise when three criteria are fulfilled: the primary insult reshapes microbial community structure; the altered microbiota is sufficient to transfer pathology or induce a host injury program independently of the initiating insult; and the host responses elicited by the dysbiotic microbiota are mechanistically distinct from those triggered by the primary insult itself. Our study fulfills these criteria and thereby provides an experimental framework for dissecting microbiota-mediated contributions to disease.

The most immediate extensions of this framework are likely to emerge within oncology and transplant medicine, where epithelial barrier injury, dysbiosis, and treatment-limiting complications frequently converge. Conditioning regimens for allogeneic hematopoietic stem cell transplantation represent a particularly compelling example, as intestinal barrier damage, microbial dysbiosis, and bloodstream infection risk are tightly linked in this setting^16,33^. Similar considerations apply to radiation-induced enteritis and proctitis, where epithelial injury and microbial disruption occur concurrently^34^, as well as to immune checkpoint inhibitor-associated colitis, a toxicity increasingly associated with microbiome composition and intestinal barrier dysfunction^32,35^. In each of these contexts, distinguishing whether dysbiosis represents a biomarker of tissue injury or a secondary effector of pathology may have direct implications for therapeutic intervention.

Our findings also raise several additional conceptual possibilities. The ability of post-chemotherapy microbiota to transfer a permeability phenotype suggests that cytotoxic therapy can leave behind a pathogenic ecological state within the gut ecosystem. Such “transmissible toxicity” raises the possibility that treatment-associated microbial configurations may propagate host injury independently of ongoing drug exposure. In addition, treatment-induced microbiome states may serve as biomarkers of susceptibility to complications such as mucositis, bacteremia, or dose-limiting gastrointestinal toxicity. Finally, the experimental framework used here may have methodological value beyond this specific context. By enabling controlled separation of microbial stimuli from direct toxicant exposure, the gut organ culture platform provides a generalizable strategy for dissecting microbiota-mediated contributions to host pathology.

Several limitations should be considered. Although the *ex vivo* gut organ culture system preserves epithelial architecture and local immune components^20,21,23–25^, it lacks systemic inputs such as circulating immune cells, endocrine signals, and long-term regenerative dynamics. Consequently, the model captures early mucosal responses to luminal perturbations but cannot fully recapitulate the systemic complexity of chemotherapy-associated disease. Future studies using gnotobiotic models colonized with defined microbial communities from Ara-C-treated donors will be necessary to identify the specific taxa or microbial metabolites responsible for barrier disruption.

In summary, our findings demonstrate that chemotherapy-induced dysbiosis is not merely a consequence of epithelial injury, but an independent driver of intestinal barrier dysfunction. By experimentally uncoupling direct drug toxicity from microbiota-mediated effects, we identify parallel and mechanistically distinct pathways through which chemotherapy destabilizes intestinal homeostasis. More broadly, these findings suggest that a component of chemotherapy toxicity is ecologically mediated and may therefore be therapeutically separable from anticancer efficacy, providing a rationale for microbiota-targeted strategies to mitigate treatment-associated injury.

## Methods

### Mice

Male and female C57BL/6J (B6) mice were obtained from Envigo RMS (Israel) and maintained under specific pathogen-free (SPF) conditions at the Bar-Ilan University animal facility. Mice were housed under controlled temperature and humidity, with a 12-h light/dark cycle and had ad libitum access to standard chow and water. Mice aged 8-12 weeks were used for *in vivo* experiments, whereas 14-day-old littermates mice were used for *ex vivo* colon organ culture experiments. All animal experiments were approved by the Bar-Ilan University Institutional Animal Care and Use Committee (IACUC) BIU-IL-21-03-2021 and conducted in accordance with relevant institutional and national guidelines for the care and use of laboratory animals.

### Cell culture

Human colorectal adenocarcinoma Caco-2 cells were cultured in Dulbecco’s modified Eagle medium (DMEM/F12) medium supplemented with 20% heat-inactivated fetal bovine serum (FBS; Gibco), 2 mM L-glutamine, and 100 U/mL penicillin-streptomycin at 37°C in a humidified incubator with 5% CO₂. For barrier measurements, cells were grown to confluent monolayers. Cells were routinely tested for mycoplasma contamination and confirmed negative.

### Gut organ culture and *ex vivo* intestinal permeability assay (X-IPA)

Gut organ culture experiments were performed as previously described^20,21,25^ to enable controlled exposure to chemotherapy and microbiota-derived stimuli. Briefly, intact colonic segments were dissected from C57BL/6 mice under sterile conditions, and luminal contents were gently flushed with pre-warmed sterile medium. Tissues were mounted onto a custom-designed gut organ culture device by securing both ends over inlet and outlet ports, preserving the native tissue architecture and luminal orientation. The culture device was maintained at 37 °C in a humidified environment and continuously perfused with oxygenated, serum-free culture medium (IMDM supplemented with 20% Knock-Out serum replacement, 2% B-27, 1% N-2, 1% L-glutamine, 1% non-essential amino acids, and 1% HEPES). A controlled gas mixture (95% O₂ / 5% CO₂) was supplied to sustain tissue viability. Luminal flow was regulated using a syringe pump to ensure stable and reproducible delivery of experimental conditions.

For permeability measurements, FITC-dextran (4 kDa, 0.5 mg mL) was infused into the intestinal lumen, either alone or in combination with the indicated stimuli. For direct drug exposure experiments, Ara-C (cytarabine) was added to the luminal perfusate at the specified concentrations. For microbiota-driven experiments, fecal samples were collected from donor mice (pre- or post-Ara-C treatment), homogenized in sterile medium (10 mg feces per 1 mL medium), strained through a 100 μm cell strainer to remove large particles, and introduced into the lumen together with FITC-dextran.

Barrier integrity was quantified by measuring the accumulation of FITC-dextran in the extraintestinal medium, which reflects trans-epithelial permeability^20^. Fluorescence in the surrounding medium was measured at defined time points (typically 2 h and 4 h) using a fluorometer (excitation 485 nm, emission 520 nm). For comparative analyses, fluorescence values were normalized to internal controls within each experiment to account for inter-sample variability. At the end of each experiment, tissues were collected for downstream analyses, including bulk RNA sequencing.

### Intraperitoneal (i.p.) Injection

Chemotherapy-induced mucositis was established in C57BL/6J mice (8-12 weeks old) by daily intraperitoneal (i.p.) injections of cytarabine (Ara-C, Sigma) at a dose of 3.6 mg per mouse, diluted in 100 μL phosphate-buffered saline (PBS) for four consecutive days ^36^. Control mice received 100 μL PBS.

### Oral gavage and *in vivo* permeability assay

For *in vivo* intestinal permeability assays, mice aged 8-12 weeks were fasted for 12 h prior to oral gavage of 150 μL containing the indicated stimuli together with 4 kDa FITC-dextran (80 mg/mL in sterile PBS) as a permeability tracer. Blood samples were collected 5 h post-gavage from the facial vein using a 25G needle, and serum fluorescence was measured using a fluorometer (excitation 485 nm, emission 520 nm).

### Fecal samples collection

Fecal samples collected mice before and after chemotherapy treatment were stored at −80 °C until use. Samples were resuspended in sterile medium or PBS under anaerobic conditions and homogenized to generate fecal bacterial suspensions.

### DNA extraction and 16S rRNA sequencing

DNA was extracted from approximately 0.25 g of fecal material using the DNeasy PowerSoil Kit (Qiagen) according to the manufacturer’s instructions. DNA concentration was measured using a NanoDrop 2000 spectrophotometer (Thermo Fisher Scientific). Negative controls were included during DNA extraction and library preparation. Libraries targeting the V4 region of the 16S rRNA gene were prepared by HyLabs (Israel) using a two-step PCR protocol, purified using KAPA Pure beads (Roche), quantified with Qubit fluorometry (Life Technologies), pooled, and size-validated using a TapeStation system. Pooled libraries were sequenced on an Illumina MiSeq platform using a MiSeq v2 kit (500 cycles).

### Analysis of 16S rRNA sequencing

To evaluate treatment-associated alterations in microbial community composition, raw paired-end FASTQ files containing 16S rRNA gene sequences were processed using the QIIME2 pipeline (v2023.7.0). Sequences were imported into QIIME2 and denoised using the DADA2 algorithm. For quality trimming, 13 bp were removed from the left of both forward and reverse reads, while forward reads were truncated at 241 bp and reverse reads at 176 bp. DADA2 was further used for quality filtering, chimera removal, and amplicon sequence variant (ASV) inference. Representative ASVs were aligned using the MAFFT program, and a phylogenetic tree was constructed with the FastTree algorithm. Taxonomic assignment was performed with the classify-sklearn naïve Bayes classifier in QIIME2 using the Greengenes2 (v2022.10). ASVs assigned to mitochondria or chloroplast were removed prior to downstream analyses.

Beta diversity analysis was performed to assess treatment-associated microbiome restructuring between matched before- and after-treatment samples. Only samples with complete paired sampling at both time points were retained for downstream analysis. The ASV count table was then rarefied to the minimum sequencing depth across all retained samples using the rrarefy() function from the vegan package in R. Each sample was subsequently normalized to relative abundance. Bray-Curtis dissimilarity matrices were computed using the vegdist() function (vegan). Principal Coordinates Analysis (PCoA) was performed using the cmdscale() function to reduce the multidimensional distance matrix into two principal axes. For clarity of visualization, a subset of representative paired samples is displayed in the PCoA plot, with paired before- and after-treatment samples connected to depict within-subject temporal microbiome trajectories, whereas statistical analysis was performed on the full paired cohort. Statistical differences in overall microbial community composition were assessed using PERMANOVA (adonis2(), 999 permutations), while constraining permutations within mouse identity (strata = MouseID) to account for the repeated-measures design.

Differential abundance analysis was conducted in R using DESeq2. Read count data were aggregated at the species taxonomic level. In the DESeq2 analysis, the design formula used was ∼ mouseID + day to account for paired repeated measurements from the same animal. Significant taxa were identified based on a p-value ≤ 0.05 and an abs(log2FC) ≥ 0.58. Volcano plots were generated in R using ggplot2.

### RNA extraction and transcriptomic analysis

Mouse colon tissues were homogenized using a bead beater, and total RNA was extracted using the RNeasy Micro Kit (Qiagen) according to the manufacturer’s instructions. RNA quality was assessed by measuring A260/280 absorbance ratios. Sequencing reads were trimmed using Trim Galore, aligned to the mouse reference genome (mm39, NCBI RefSeq) using STAR (v2.7.10b), and gene counts were generated using HTSeq-count (v2.0.4).

Differential gene expression analysis was performed in R using DESeq2. In all RNA-seq comparisons, the DESeq2 design formula used was ∼ gender + treatment in order to account for sex-associated transcriptional variability while testing for treatment-dependent effects. Genes were considered significantly differentially expressed if they met the thresholds of *P*-value ≤ 0.05 and |log2 fold change| ≥ 0.58 (corresponding to a fold change of ≥1.5).

For pathway enrichment analysis, significantly upregulated and downregulated gene lists were exported separately and analyzed using Metascape. Differential expression visualizations, including volcano plots and log2 fold change cross-comparison scatter plots, were generated in R using ggplot2 based on the DESeq2 output.

### TEER measurements

Caco-2 epithelial cells were seeded onto 96-well CytoView-Z impedance plates (Axion BioSystems; cat. Z96-IMP-96B-25) at a density of 5 × 10⁵ cells per well and maintained under standard culture conditions until reaching full confluence (800-1200 Ω). TEER measurements were performed using the Maestro Edge platform (Axion BioSystems) at 37 °C with 5% CO₂, and data were acquired using AxIS Z software (v3.2.3).

Barrier integrity was quantified as the barrier index, calculated by the Axion impedance module as the ratio between cellular resistance at low frequency (1 kHz) and high frequency (41 kHz). Measurements were recorded at 1-minute intervals for up to 24 hours and normalized to the baseline value (t = 0). Where indicated, additional normalization was performed relative to unstimulated control wells.

### HPLC analysis

HPLC analysis was performed at the BIU Faculty of Life Sciences core facility. Sample preparation method for High-performance liquid chromatography) HPLC (analysis was adapted from procedure of Y. Sun et al. J. Chromatography B, 870 (2008). Solid-phase extraction was done using 1mL Oasis MCX cartridge (Waters Corp). The fecal specimens were resuspended in 1mL double distilled water and homogenized using a bead beater (Tissue Lyser II, Qiagen) according to the manufacturer protocol. The supernatant was transferred to the solid-phase extraction cartridge preconditioned with 1mL methanol and 1mL of water. The sample was washed twice with 1mL of 0.05M HCl and 5% methanol, followed by 1mL of methanol. Elution of Ara-C was accomplished by adding methanol containing ammonium hydroxide (95:5 v/v). The samples were evaporated using N_2_ sample concentrator device (Techne FSC496D). Dried samples were reconstituted with 150 μL of water.

The chromatography apparatus consisted of Hitachi Elite LaChrom system equipped with diode array detector, column oven, autosampler, and a quaternary pump. All chromatographic analyses were performed at 30 °C using HIChrom RPB (5 μm particle size, L × I.D. 25 cm × 4.6 mm), flow rate 1.0 mL/min under isocratic elution conditions with the following buffer composition: 25 mM potassium phosphate, pH 3.2 (adjusted with phosphoric acid): Acetonitrile (99:1). Each analysis cycle was set to 15 minutes with injection volume of 5 μL. The chromatographic flow was monitored at 260 nm and integrated using EZChrom Elite Software.

### Quantification and statistical analysis

Statistical analyses were performed using GraphPad Prism (version 10(. Statistical parameters, including the definition and exact value of n, are reported in the corresponding figure legends. Comparisons between two groups were performed using two-tailed Student’s t-tests, while multiple group comparisons were analyzed using one-way ANOVA with appropriate post hoc tests, as indicated in the figure legends. Differences were considered statistically significant at p ≤ 0.05.

## Supporting information

Summary Supplemental Table

## Acknowledgements

We thank Dr. Alexander Varvak for HPLC analysis and members of the Yissachar lab for insightful discussions. This work was supported by the Israel Science Foundation (grant no.1384/18, 2346/25, N.Y.) and the Israel Cancer Association (grant no.20191634, N.Y.).

