## Supplementary Figures for "Uncoupling Drug and Microbiota Contributions to Chemotherapy-Induced Gut Toxicity"

**Fig. S1**

**a,** Metascape enrichment analysis of DEGs in intestinal tissues. The analysis compares 100mM Ara-C versus 10mM Ara-C concentrations following a 2 h stimulation period. Significantly downregulated biological pathways are ranked by significance ($-\log_{10}P$).

**b-c,** Metascape enrichment analysis of DEGs in intestinal tissues. The analysis compares 100mM Ara-C concentration versus control (sterile medium) following a 2 h stimulation period. Bars represent significantly **b** upregulated and **c** downregulated biological pathways, ranked by significance ($-\log_{10}P$).

**d-e,** Metascape enrichment analysis of DEGs in intestinal tissues. The analysis compares 10mM Ara-C concentration versus control (sterile medium) following a 2 h stimulation period. Bars represent significantly **d** upregulated and **e** downregulated biological pathways, ranked by significance ($-\log_{10}P$).

**Fig. S2 Ara-C treatment alters host physiology *in vivo*.**

**a,b,** Body weight dynamics of female (a) and male (b) mice treated with Ara-C or PBS. Left panels show weight change at day 5 as a percentage of initial body weight. Right panels show longitudinal body weight changes from day 1 to day 5. (female: PBS n = 24, Ara-C n = 23; male: PBS n = 12, Ara-C n = 41).

**c,** Colon length of control (PBS, n = 22) and Ara-C-treated mice (n = 35) measured at day 5. No significant difference was observed (ns).

**Statistics.** Statistical analyses were performed using t-test, as indicated in the respective panels. Data are presented as mean ± s.d. Significance thresholds: ***P < 0.0001, ***P < 0.001, **P < 0.01, *P < 0.05; ns, not significant.

**Fig. S3**

**a,** Metascape enrichment analysis of DEGs in intestinal tissues. The analysis compares responses to post- versus pre-Ara-C microbiota following a 4 h stimulation period. Bars represent significantly downregulated biological pathways, ranked by significance ($-\log_{10}P$).

**b-c,** Metascape enrichment analysis of DEGs in intestinal tissues. The analysis compares responses to post- versus pre-Ara-C microbiota following a 2 h stimulation period. Bars represent significantly **b** upregulated and **c** downregulated biological pathways, ranked by significance ($-\log_{10}P$).

Fig. S1

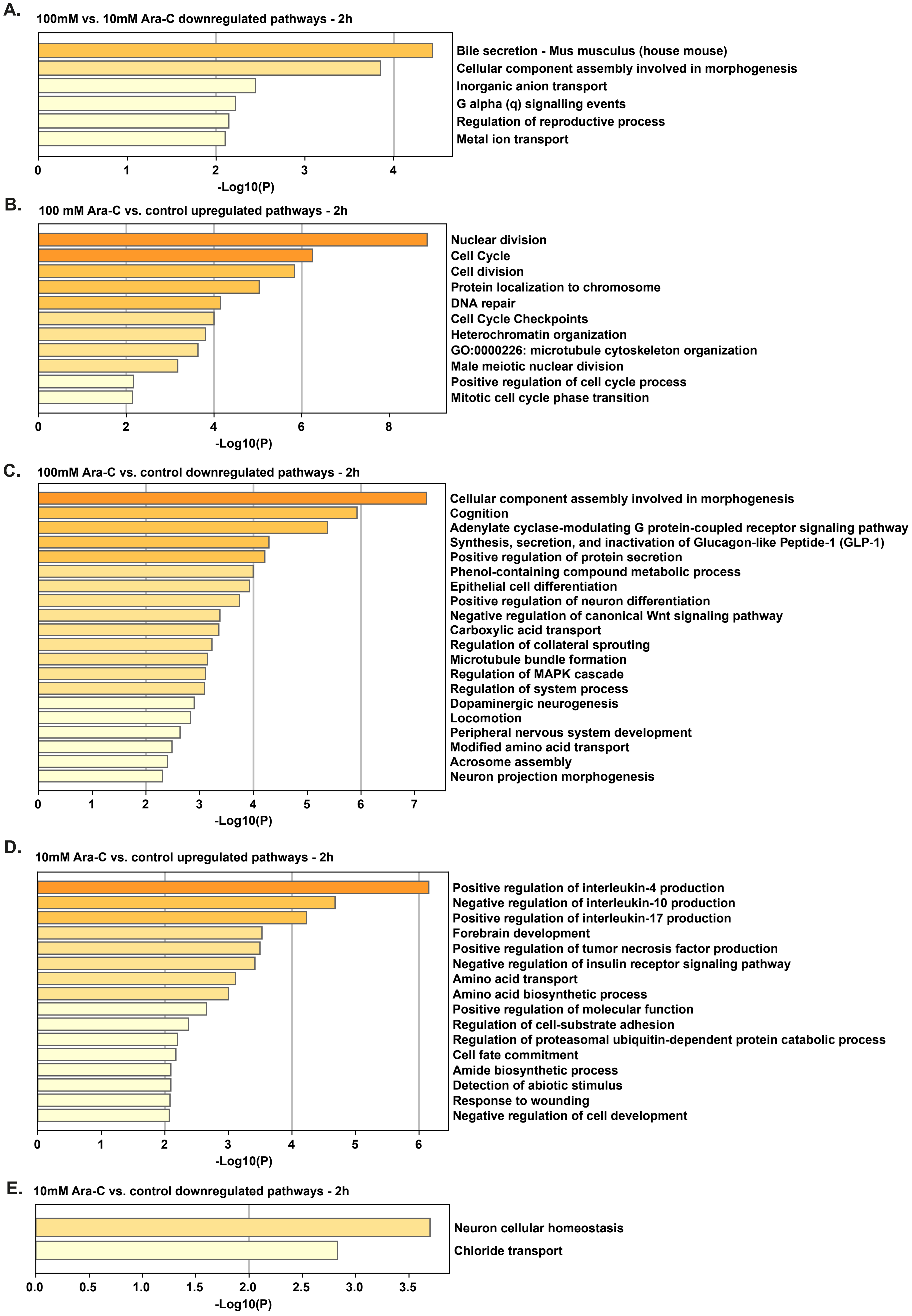
A.
100mM vs. 10mM Ara-C downregulated pathways - 2h
Bile secretion - Mus musculus (house mouse)
Cellular component assembly involved in morphogenesis
Inorganic anion transport
G alpha (q) signalling events
Regulation of reproductive process
Metal ion transport
0
1
2
3
4
-Log10(P)
B.
100 mM Ara-C vs. control upregulated pathways - 2h
Nuclear division
Cell Cycle
Cell division
Protein localization to chromosome
DNA repair
Cell Cycle Checkpoints
Heterochromatin organization
GO:0000226: microtubule cytoskeleton organization
Male meiotic nuclear division
Positive regulation of cell cycle process
Mitotic cell cycle phase transition
0
2
4
6
8
-Log10(P)
C.
100mM Ara-C vs. control downregulated pathways - 2h
Cellular component assembly involved in morphogenesis
Cognition
Adenylate cyclase-modulating G protein-coupled receptor signaling pathway
Synthesis, secretion, and inactivation of Glucagon-like Peptide-1 (GLP-1)
Positive regulation of protein secretion
Phenol-containing compound metabolic process
Epithelial cell differentiation
Positive regulation of neuron differentiation
Negative regulation of canonical Wnt signaling pathway
Carboxylic acid transport
Regulation of collateral sprouting
Microtubule bundle formation
Regulation of MAPK cascade
Regulation of system process
Dopaminergic neurogenesis
Locomotion
Peripheral nervous system development
Modified amino acid transport
Acrosome assembly
Neuron projection morphogenesis
0
1
2
3
4
5
6
7
-Log10(P)
D.
10mM Ara-C vs. control upregulated pathways - 2h
Positive regulation of interleukin-4 production
Negative regulation of interleukin-10 production
Positive regulation of interleukin-17 production
Forebrain development
Positive regulation of tumor necrosis factor production
Negative regulation of insulin receptor signaling pathway
Amino acid transport
Amino acid biosynthetic process
Positive regulation of molecular function
Regulation of cell-substrate adhesion
Regulation of proteasomal ubiquitin-dependent protein catabolic process
Cell fate commitment
Amide biosynthetic process
Detection of abiotic stimulus
Response to wounding
Negative regulation of cell development
0
1
2
3
4
5
6
-Log10(P)
E.
10mM Ara-C vs. control downregulated pathways - 2h
Neuron cellular homeostasis
Chloride transport
0.0
0.5
1.0
1.5
2.0
2.5
3.0
3.5
-Log10(P)

Fig. S2

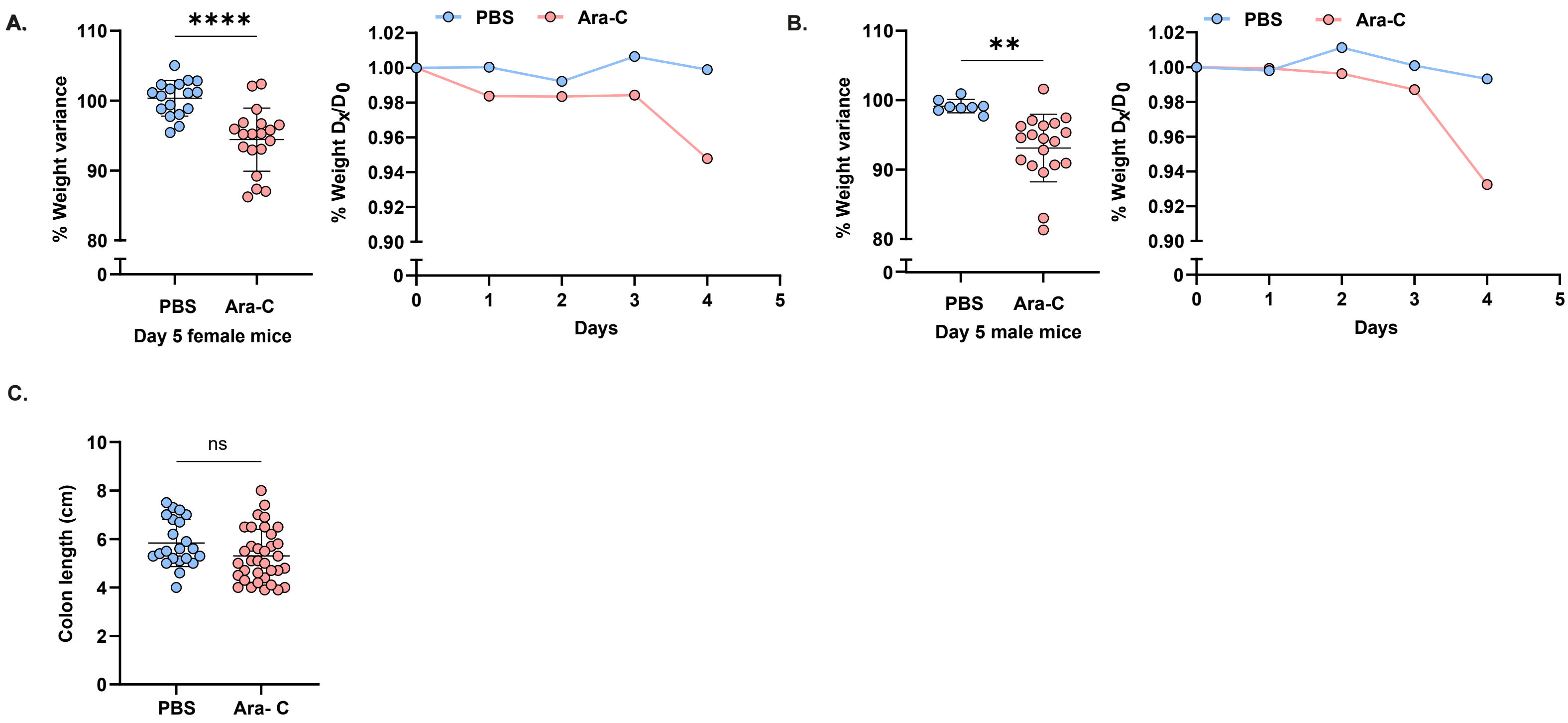
A.
****
% Weight variance
110
100
90
80
0
PBS
Ara-C
Day 5 female mice
PBS
Ara-C
% Weight Dx/D0
1.02
1.00
0.98
0.96
0.94
0.92
0.90
0
1
2
3
4
5
Days
B.
**
% Weight variance
PBS
Ara-C
Day 5 male mice
PBS
Ara-C
% Weight Dx/D0
Days
C.
ns
Colon length (cm)
10
8
6
4
2
0
PBS
Ara- C

### Fig. S3

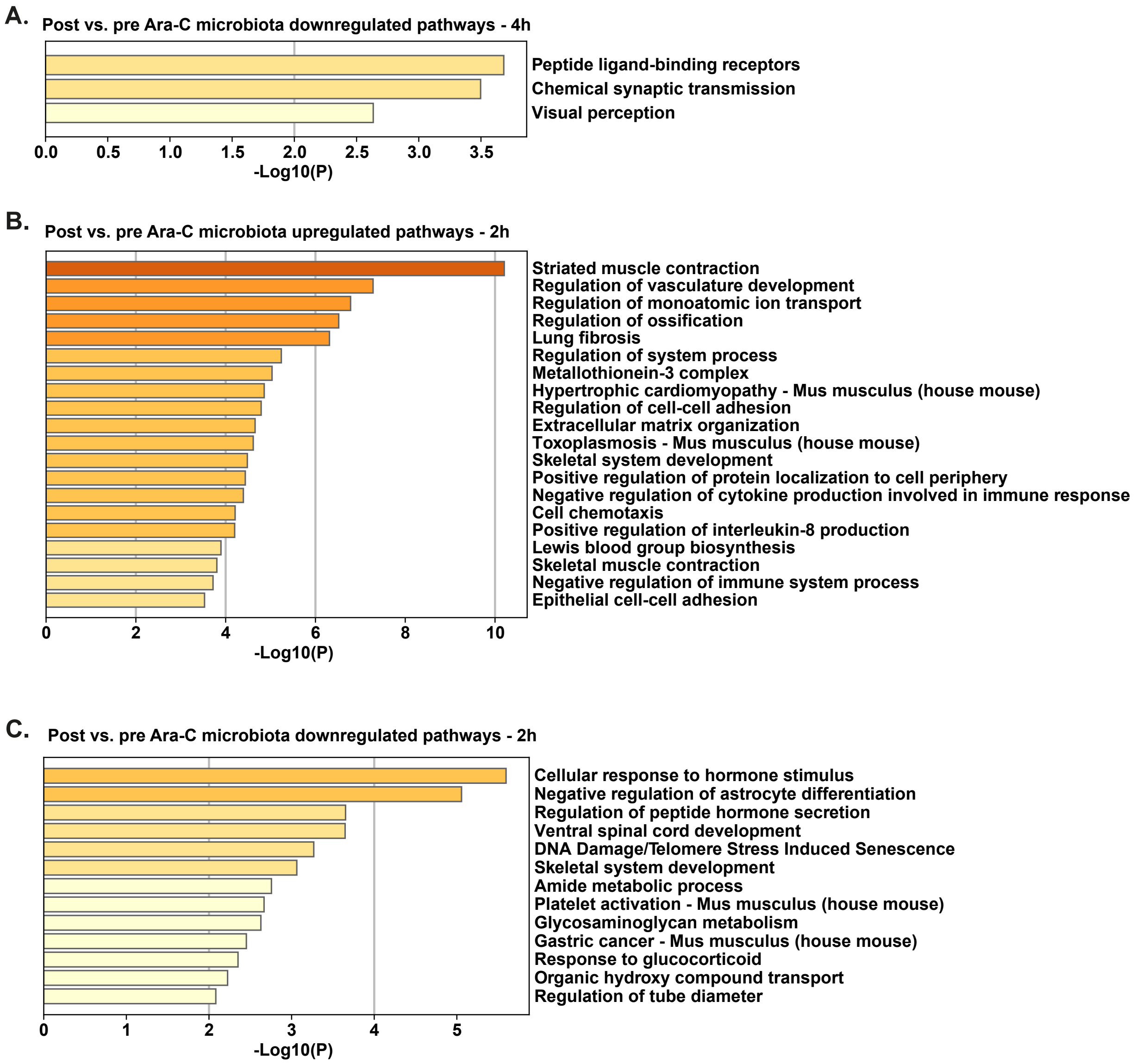
A.
Post vs. pre Ara-C microbiota downregulated pathways - 4h
Peptide ligand-binding receptors
Chemical synaptic transmission
Visual perception
0.0 0.5 1.0 1.5 2.0 2.5 3.0 3.5
-Log10(P)
B.
Post vs. pre Ara-C microbiota upregulated pathways - 2h
Striated muscle contraction
Regulation of vasculature development
Regulation of monoatomic ion transport
Regulation of ossification
Lung fibrosis
Regulation of system process
Metallothionein-3 complex
Hypertrophic cardiomyopathy - Mus musculus (house mouse)
Regulation of cell-cell adhesion
Extracellular matrix organization
Toxoplasmosis - Mus musculus (house mouse)
Skeletal system development
Positive regulation of protein localization to cell periphery
Negative regulation of cytokine production involved in immune response
Cell chemotaxis
Positive regulation of interleukin-8 production
Lewis blood group biosynthesis
Skeletal muscle contraction
Negative regulation of immune system process
Epithelial cell-cell adhesion
0 2 4 6 8 10
-Log10(P)
C.
Post vs. pre Ara-C microbiota downregulated pathways - 2h
Cellular response to hormone stimulus
Negative regulation of astrocyte differentiation
Regulation of peptide hormone secretion
Ventral spinal cord development
DNA Damage/Telomere Stress Induced Senescence
Skeletal system development
Amide metabolic process
Platelet activation - Mus musculus (house mouse)
Glycosaminoglycan metabolism
Gastric cancer - Mus musculus (house mouse)
Response to glucocorticoid
Organic hydroxy compound transport
Regulation of tube diameter
0 1 2 3 4 5
-Log10(P)
